# *detectEVE*: fast, sensitive and precise detection of endogenous viral elements in genomic data

**DOI:** 10.1101/2024.09.06.611620

**Authors:** Nadja Brait, Thomas Hackl, Sebastian Lequime

**Affiliations:** Cluster of Microbial Ecology, Groningen Institute for Evolutionary Life Sciences, University of Groningen, Groningen, The Netherlands

## Abstract

**Summary:** Endogenous viral elements (EVEs) are fragments of viral genomic material embedded within the host genome. Retroviruses contribute to the majority of EVEs due to their genomic integration during their life cycle, however, the latter can also arise from non-retroviral RNA or DNA viruses, then collectively known as non-retroviral (nr)EVEs. Detecting nrEVEs poses challenges due to their sequence and genomic structural diversity, contributing to the scarcity of specific tools designed for nrEVEs detection.

Here, we introduce *detectEVE*, a user-friendly and open-source tool designed for the accurate identification of nrEVEs in genomic assemblies. *detectEVE* deviates from other nrEVE detection pipelines, which usually classify sequences in a more rigid manner as either virus-associated or not. Instead, we implemented a scaling system assigning confidence scores to hits in protein sequence similarity searches, using bit score distributions and search hints related to various viral characteristics, allowing for higher sensitivity and specificity. Our benchmarking shows that *detectEVE* is computationally efficient and accurate, as well as considerably faster than existing approaches, due to its resource-efficient parallel execution.

Our tool can help to fill current gaps in both host-associated fields and virus-related studies. This includes (i) enhancing genome annotations with metadata for EVE loci, (ii) conducting large-scale paleo-virological studies to explore deep viral evolutionary histories, and (iii) aiding in the identification of actively expressed EVEs in transcriptomic data, reducing the risk of misinterpretations between exogenous viruses and EVEs.

**Availability and Implementation:** *detectEVE* is implemented as snakemake workflow, available with detailed documentation at https://github.com/thackl/detectEVE and can be easily installed using conda.

## Introduction

Endogenous viral elements (EVEs) arise from the integration of viral genomic material into the germline cells of the host genome, which can then be inherited by its progeny [1,2]. EVEs can provide new functions to their hosts [3], but can also be seen as viral fossils, providing insights into the evolutionary history of host-virus interactions and host-viral defense mechanisms [4,5]. EVEs can be categorized into two main groups: endogenous retroviruses (ERVs) and non-retroviral endogenous viral elements (nrEVEs). While nrEVEs are less common than ERVs, representatives have been found across all eukaryotic groups [6–10].

Thanks to advances in sequencing technologies and associated computational methods, the quantity of sequenced genomes is increasing rapidly. As of September 2024, over 2.3 million genome assemblies were available on NCBI, including over 42.7K eukaryotic assemblies [11]. This abundance of data provides a rich resource for gaining new insights into ERVs and nrEVEs but also creates the need for computationally efficient analysis workflows.

Most available tools are built for ERV or TE detection or the classification of EVEs derived from specific viral families [12]. nrEVEs lack distinguished biological characteristics, and their high viral diversity and partial genomic integration make their identification difficult. Bioinformatic workflows for nrEVE identification in genomic data are primarily based on homology-based methods, such as BLAST+ [13], DIAMOND [14], or HMM searches [15], and, although multiple identification studies have been conducted [16–20], ready-to-use pipelines are lacking. One of the few tools available for nrEVE detection is the Database Integrated Genome Screening (DIGS) tool [21,22]. Its efficient use, however, requires experience in EVE detection, and it has yet to be used outside of the group that developed it.

Here, we present *detectEVE*, a user-friendly, open-source tool for detecting and annotating nrEVEs in genome assemblies. In addition to sequence similarity searches, our tool features an EVE scoring algorithm that considers bitscore distributions and sequence hit origins to classify and annotate potential EVE sequences into confidence categories. Our benchmarking demonstrates excellent performance and detection sensitivity compared to other EVE detection tools.

### The *detectEVE* framework

*detectEVE* is a fully automated, command-line tool for detecting non-retroviral endogenous viral elements (nrEVEs) in host genomes. The workflow is organized into four steps: (I) identification of putative nrEVEs (putatEVEs) from viral input sequences in genomic assemblies through sequence similarities, (II) cleaning steps, (III) exclusion of false positives through a second sequence similarity search against a general protein database, and (IV) applying an EVE scoring system for the categorization of validated EVEs (validatEVEs) into confidence levels (Figure 1).

**Figure 1.**
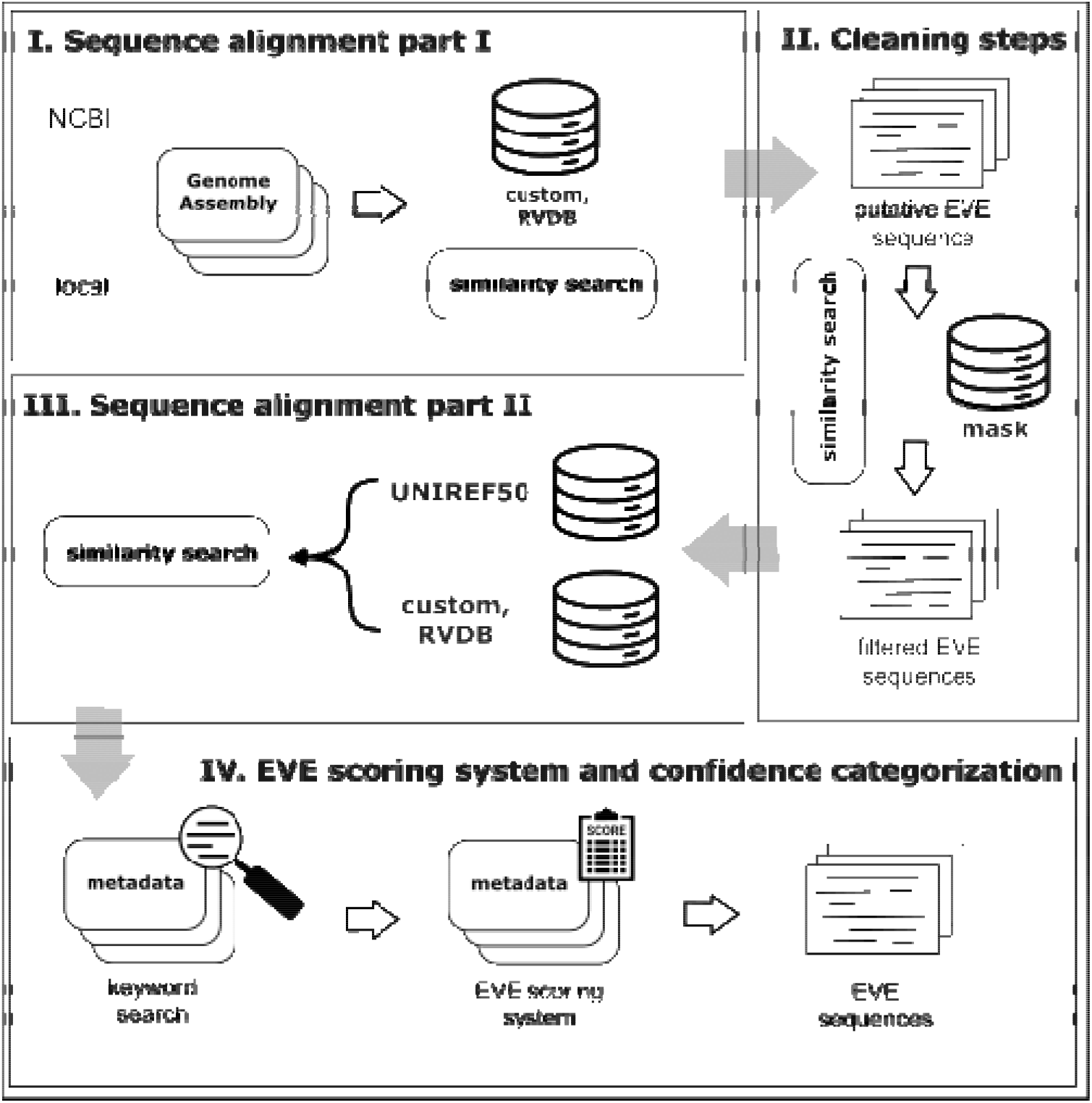
The *detectEVE* framework -a schematic view.

*detectEVE* works either on user-provided genomes in fasta format or on a list of genome accessions from NCBI/WGS/Traces (https://www.ncbi.nlm.nih.gov/Traces/wgs/), for which assemblies will be automatically downloaded. putatEVEs are identified by comparing genomes against a virus database with DIAMOND [14]. By default, the genomes are searched against a clustered version of the Reference Viral Databases (RVDB-80) [23], a proteic database of (non-bacterial) viruses and viral-like sequences, frequently updated from NCBI sequence databases. *detectEVE*’s default search is restricted to *Orthornavirae* (non-retroviral RNA viruses -taxid:2732396) and *Monodnaviria* (ssDNA -taxid:2731342). However, custom taxon searches, as well as custom input databases, are supported. putativEVEs resembling known false positive sequences are masked based on a customizable masking database.

To assess the validity of putatEVEs, we match them back against two databases: (I) RVDB-80 ± input viral database and (II) a clustered UNIREF-50 [24] database. Matching to both databases enhances specificity, as UNIREF filters false positives, while RVDB and input databases retain viral hits that UNIREF might miss, resulting in a more accurate hit distribution. The combined hits are filtered with hits below a default threshold of 50% of the top score being removed (*B*_min_=*B*_top_×0.5). Further on, *detectEVE* integrates descriptive information of each hit quantitatively, allowing it to rank and assess hit quality easily. To do so, we devised a system that scores EVEs on a scale from 0 to 100, representing an overall measure of the likelihood of identifying an EVE. To obtain these scores, we first categorize hits into categories using one of the following tags: “viral”, “maybe viral”, “retroviral”, “non viral” and “false viral”, corresponding to assigned keywords matching words in the description of each hit (see Supplementary Table 1). The tag “maybe viral” is used for uncharacterized or hypothetical proteins in host genomes, as these may be EVEs with intact open reading frames. Then, we aggregate scores according to the formula

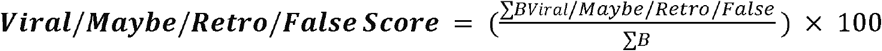

with ***B*** being the total bitscore for all hits for that specific locus. These individual scores allow the calculation of our total EVE score with the formula:

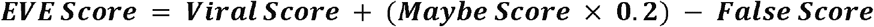

Based on these scores, *detectEVE* classifies each validatEVE into one of three quality tiers -high confidence (EVE score >30), low confidence (EVE score: 10–29), or no confidence (< 10, excluded from the final output). Since *detectEVE* is not optimized to differentiate between ERVs or retrotransposons, hits with a Retro score of 10 or above are automatically given low confidence. Finally, *detectEVE* produces three output files: a ‘<genome_id>-validatEVEs.fna’ file containing all EVE nucleotide sequences, a ‘<genome_id>-validatEVEs.pdf’ file providing a graphical overview of hit distributions, and a ‘<genome_id>-validatEVEs.tsv’ file. The TSV file includes confidence levels, EVE-score calculations, and metadata such as EVE locus, top e-value, top percent identity, top description, top viral description, and viral taxonomy parameters. These files ensure optimal data accessibility for functional annotations and facilitate downstream analyses.

*detectEVE* is built on Snakemake, a Python-based, user-friendly, fast, and intuitive workflow management system [25]. Snakemake also effectively executes tasks in parallel, allowing detectEVE to scale well on large datasets. All parameters can be easily adjusted to the user’s needs in a central configuration file, allowing users to customize their workflows without modifying the underlying code. Using conda for dependency management, *detectEVE* can be easily installed using the provided environment configuration. Default databases can be readily downloaded and set up using a dedicated command.

We have specifically chosen to exclude viral dsDNA virus-derived and retrovirus-derived EVEs from the default analysis in *detectEVE*. While it is theoretically possible to use *detectEVE* for these detections, the differentiation between partial sequences of exogenous dsDNA viruses in genome assemblies and viral integrations remains challenging. Several specialized approaches have been developed for the identification of prophages in complete microbial genomes (e.g., geNomad [26], VIBRANT [27], PHASTEST [28]) or retroviral integrations (e.g., RepeatMasker [29], RetroSeq [30], or ERVcaller [31]). *detectEVE*, however, provides an open framework for future developments, which can include detecting all EVE types.

### Performance evaluation and comparison to other tools

We compared *detectEVE’s* performance to other bioinformatic methods developed for EVE detection, namely 1) DIGS -A Database-Integrated Genome Screening Tool, which applies a similarity search-based screening pipeline linked to a relational database management system [22]; and 2) a protocol developed by Palatini et al., for the discovery and annotation of viral integrations [18]. DIGS and Palatini’s protocol were installed according to their installation guides.

We first estimated speed and CPU performance based on a single input genome (*Anopheles gambiae*: AAAB01 -328.94 Mbp) utilizing 1, 8, 16, and 40 threads on an Intel(R) Xeon(R) Gold 5215 CPU @ 2.50GHz processor. *detectEVE* has great multi-threading efficiency, i.e., the total run time decreases clearly with the number of threads used, which shows efficient CPU usage vs. time (Supplementary Table 2, Supplementary Figure 1). In contrast, DIGS’s or Palatini’s multi-threading shows no significant reduction in runtime beyond 8 threads and 16 threads, respectively, taking nearly 42 times and 31 times longer to screen compared to *detectEVE*.

We evaluated the sensitivity and specificity of *detectEVE* by replicating results from a previous study on *Orthoflavivirus*-derived EVEs in *Anopheles* mosquito genomes [32]. To ensure fair comparisons, we used the same input data for all three applications: 24 *Anopheles* genome assemblies (accessions in Supplementary Table 3) and a custom viral database of Flavivirus accessions (NCBI:txid11050) filtered from *detectEVE’s* clustered RVDB database, containing a total of 856 sequences. Since the other applications do not support automated genome downloads, sequence files were provided manually. Runtimes were approximately 2.5 hours for *detectEVE*, 17 hours for DIGS, and 25 hours for the Palatini protocol (see Supplementary table 2). The original 2017 study identified three EVEs, all detected by every tool, with *detectEVE* classifying them as high-confidence hits (Supplementary Figure 2, Supplementary table 4). Furthermore, an updated *Flavivirus* reference list enabled all tools to detect one additional EVE. Moreover, *detectEVE* identified two more EVEs of high confidence, confirmed through manual BLASTx analysis. As DIGS lacks an automatic exclusion of false positives through a second sequence similarity search, the tool identified 347 potential EVE hits, most of which matched the Bovine viral diarrhea virus (Pestivirus), which is a known false positive [33] (Supplementary table 5).

Sequence lengths of detected EVEs varied slightly between all tools (± 1%-17% discrepancy) (Supplementary Figure 2). These discrepancies arise from the pipeline algorithms: DIGS and the Palatini protocol use the BLAST+ suite [13], while detectEVE employs DIAMOND [14] for initial sequence similarity searches, with differing parameters such as word size and gap penalties. Additional factors like range-culling, frameshift detection, and sensitivity settings also influence sequence length. Notably, longer sequences do not inherently indicate superior alignment or accuracy, as shorter sequences can sometimes provide more precise coverage.

### Tool validation on a broad viral and host spectra

To assess the sensitivity of *detectEVE* on a broader scale, we focused our efforts on detecting EVEs from four major virus groups: ssDNA viruses, negative-strand nrRNA viruses, positive-strand nrRNA viruses, and dsRNA viruses in a broad spectrum of hosts. We benchmarked *detectEVE* against data from four different hallmark papers, which looked at 25 genomes from animals, plants, and fungi [34–37] and identified and described 190 EVEs(see Supplementary table 6 and Supplementary table 7). We used the clustered RVDB database for this benchmark as a viral reference input. To avoid bias in EVE detection, both the RVDB and UNIREF50 databases were checked for the presence of sequences submitted by the authors of these studies. As some of the target EVEs showed low amino-acid similarity scores to viruses in the original manuscripts (below 30%), DIAMOND sensitivity settings in *detectEVE* were adjusted to ‘very-sensitive’.

*detectEVE* was able to detect 185 out of 190 described EVEs (sensitivity > 97%) (see Supplementary table 8), of which 174 passed *detectEVE’s* confidence thresholds (Supplementary table 9). Out of these, 167 EVEs (96 %) received a high confidence score, while only 7 were classified as low confidence. Ten out of eleven non-passing EVEs had sufficient viral-origin hits, however, the highest bitscore values corresponded to annotated host proteins lacking descriptors for viral or endogenous origin (Supplementary tables 10 and 11). Finally, one previously described EVE did not exceed EVE-score thresholds due to its high sequence similarity to myosin-9. Only one of the five EVEs that could not be detected at all with *detectEVE* was detected with a web-based blastx search.

## Conclusion

*detectEVE* is an open-source Snakemake tool for the fast, sensitive, and precise detection of EVEs in genomic assemblies, offering minimal installation requirements, high flexibility, and user-friendliness through comprehensive documentation. Our benchmarking shows that *detectEVE* is computationally efficient and accurate and considerably faster than existing approaches due to its resource-efficient parallel execution. Adjustable sensitivity settings allow *detectEVE* to be used for both paleo-virological studies, enabling extensive EVE searches to explore deep viral evolutionary histories and large-scale screenings for host genome annotations [38]. The tool can further be used to aid in the discovery of transcribed EVEs within RNA-Seq datasets by identifying EVEs in associated genomic assemblies [39]. This makes *detectEVE* applicable to a broad range of scientific applications to address current gaps in both host-associated and virus-related studies.

## Competing interests

No competing interest is to be declared.

Acknowledgments

We thank the Center for Information Technology of the University of Groningen for their support and for providing access to the Peregrine and Hábrók high-performance computing cluster.

## Data Availability

*detectEVE* is available at https://github.com/thackl/detectEVE, along with detailed documentation on installation, parameters, and FAQs (https://github.com/thackl/detectEVE/wiki). Supplementary files can be additionally found on Figshare at (Link will be provided upon acceptance).

## Author contributions

NB, TH, and SL designed the study. NB and TH implemented the software and wrote the documentation. NB carried out the benchmarking and evaluation with contributions from SL. NB, TH, and SL wrote the manuscript. TH and SL supervised the project.

## Supplementary material

### Supplementary tables

Supplementary table 1: keywords for tag classification

Supplementary table 2: Summary values and individual values of tool performances

Supplementary table 3: Anopheles genome accessions used in this study

Supplementary table 4: EVE detection for tools *detectEVE*, DIGS and Palatini

Supplementary table 5: UNIREF search against DIG EVEs for false positive detection

Supplementary table 6: description of target EVEs from hallmark papers

Supplementary table 7: accessions for genomes from hallmark papers used in this study

Supplementary table 8: detected EVEs for sensitivity estimations

Supplementary table 9: detected EVEs passing *detectEVE’*s confidence thresholds

Supplementary table 10: undetected and discarded EVEs

Supplementary table 11: hits and bitscore distributions of discarded EVEs

### Supplementary figures

**Supplementary Figure 1:**
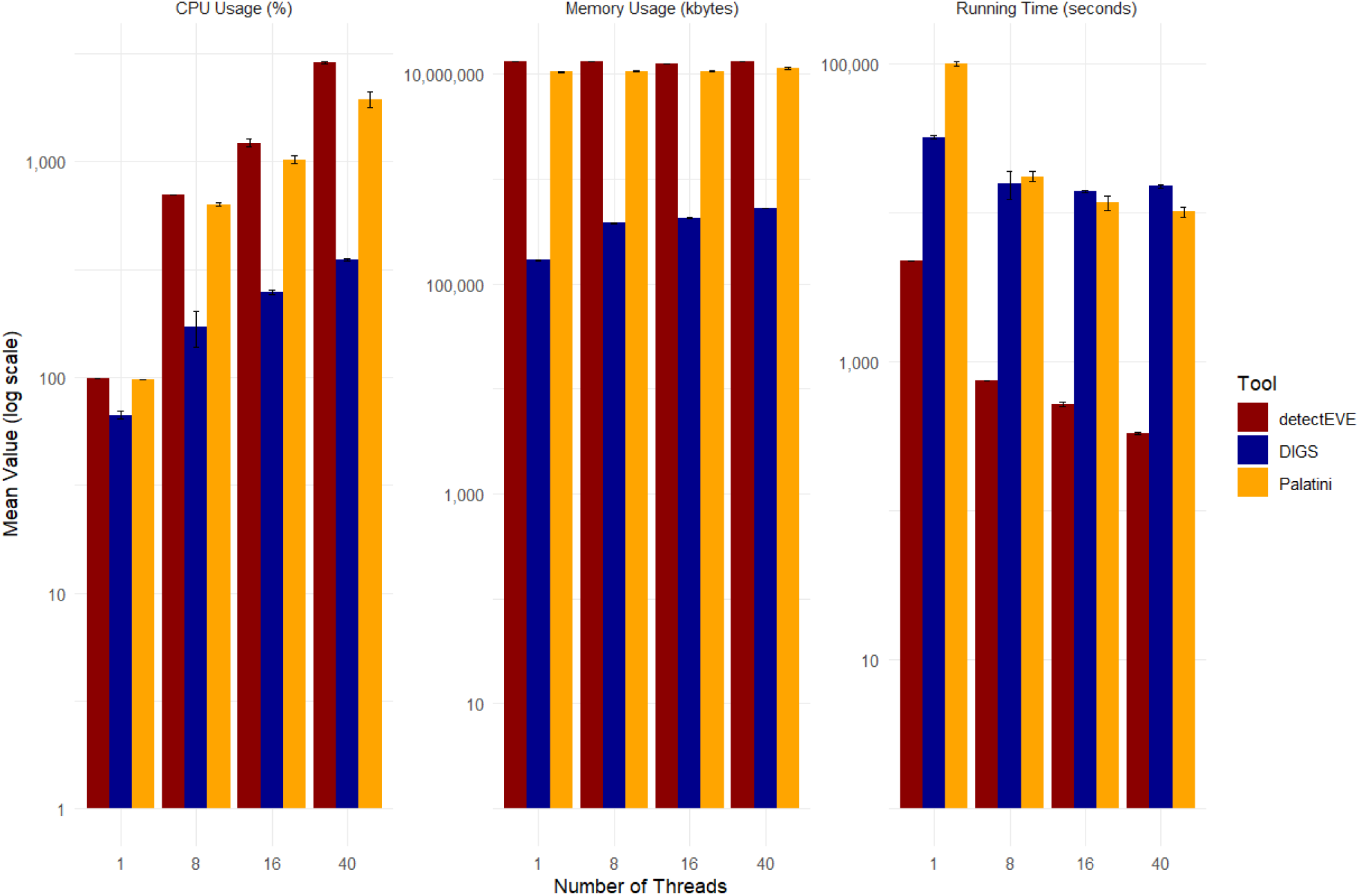
Comparison of tool performances with respect to CPU Usage, Memory Usage and running time. Analysis of sample “AAAB01” was performed in triplicates for 1, 8, 16 and 40 threads for the tools detectEVE, DIGS and the Palatini pipeline (Supplementary table 2). Mean values and standard deviations were calculated and metrics were visualized in ggplot2.

**Supplementary Figure 2:**
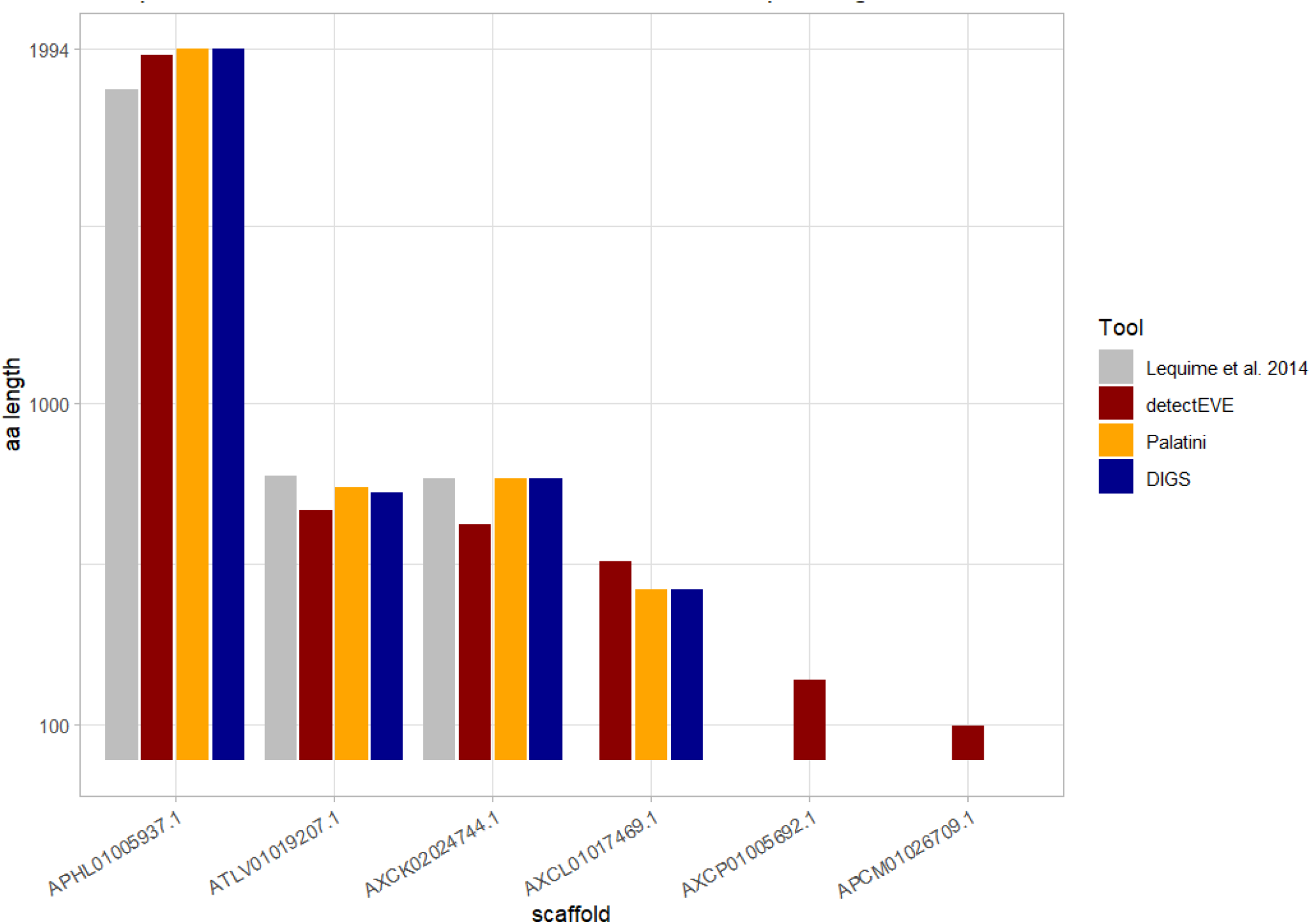
Tool comparison of presence and length of detected Flavirus EVEs in *Anopheles* genomes. 24 Anopheles genomes were screened with default settings with the tools detectEVE, DIGS, and the Palatini pipeline and compared against EVEs from Lequime et al., 2017. Presence and aa lengths of EVEs are visualized in ggplot2.

